# Conformational entropy of intrinsically disordered proteins bars intruders from biomolecular condensates

**DOI:** 10.1101/2023.03.03.531005

**Authors:** Vladimir Grigorev, Ned S. Wingreen, Yaojun Zhang

## Abstract

It has recently been discovered that eukaryotic cells are host to a multiplicity of biomolecular condensates. These condensates typically contain protein and/or RNA components with intrinsically disordered regions (IDRs). While IDRs have been proposed and demonstrated to play many roles in condensate biology, we suggest here an additional crucial role of IDRs, which is to exclude unwanted “intruders” from condensates. This exclusion effect arises from the large conformational entropy of IDRs, i.e., there is a high free-energy cost to occupying space that would otherwise be available to the IDRs. By combining polymer theory with sticker-spacer simulations, we show that the relevant insertion free energy increases with the concentration of IDRs in the condensate as well as with intruder size, attaining a linear scaling with surface area for large intruders. We find that at realistic IDR concentrations, particles as small as the size of an average protein (4 nm in diameter) can be more than 97% excluded from condensates. To overcome this entropic barrier, molecules must interact favorably with condensate components to be recruited as clients into condensates. Application of the developed size-exclusion theory to biological condensates suggests that condensate IDRs may play a generic exclusionary role across organisms and types of condensates.

---

Many of the macromolecular components of cells – proteins, RNAs, and DNA – are now known to be spatially partitioned into distinct phase-separated condensates [1, 2]. However, what leads particular components to be included in or excluded from each type of condensate is still not well understood. The components of biomolecular condensates are generally divided into two categories, scaffolds and clients [3]: The small number of components responsible for condensate formation are referred to as scaffolds. Deletion or depletion of these molecules can abolish phase separation. Other molecules that are located within the structure but do not drive phase separation are referred to as clients. Clients are typically recruited into a condensate through binding to scaffolds. It is however not clear how condensates keep out irrelevant molecules. Specifically, for an intracellular condensate immersed in a cellular milieu, it faces thousands of different types of molecules. How does it exclude unwanted molecules to maintain a specific set of desired components?

A notable feature of many of the scaffold proteins and RNAs that drive phase separation is that they contain large intrinsically disordered regions (IDRs). These IDRs are believed to play many roles, including modulating condensate stability, modifying condensate physical properties such as density and internal diffusivity, and providing binding sites for clients [7]. We propose here an additional important role of IDRs which is to exclude large undesired macromolecules from condensates (Fig. 1). Indeed, it has been observed that condensates follow a generic size-exclusion principle, i.e., they exclude large molecules. For example, while 10 kDa dextran molecules strongly partition into LAF-1 and NPM1 droplets *in vitro* and *in vivo*, 70 and 155 kDa dextran molecules are mostly excluded [8]. Similarly, perinuclear P granules in *Caenorhabditis elegans* exclude molecules larger than ∼ 70 kDa [9]. Another example is the algal pyrenoid which excludes proteins larger than ∼ 80 kDa [10]. Recently, Treen *et al*. [11] reported the discovery of repressive condensates in *Ciona* embryos, and suggested that these condensates might repress transcription by excluding the large transcriptional machinery, including RNA polymerases (∼ 500 kDa), Mediator complexes (∼ 1 MDa), and transcriptionally activating histone modifiers and nucleosome remodelers (∼ 1 *−* 3 MDa). In addition, Galagedera *et al*. [12] combined experiment and theory to show that there is a significant inclusion free-energy penalty ∼ 10 *k*_B_*T* to introduce polyubiquitin hubs into UBQLN2 condensates, and this inclusion energy increases quickly with increasing hub size. Moreover, nuclear pore complexes are known to exclude proteins above a typical size cutoff of 40 kD (∼ 5 nm in diameter) [13, 14]. Notably, all these systems contain polymers with large IDRs.

**FIG. 1.**
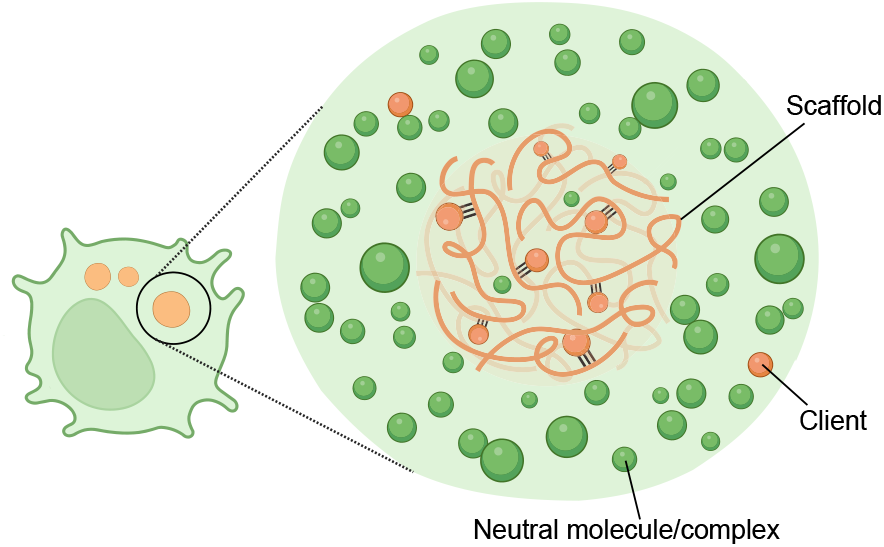
Schematic of particle exclusion and recruitment by a biomolecular condensate, prepared using BioRender [4].

Due to the polymer nature of IDRs, if the scaffolds span the entire condensate and form a stable gel network with fixed mesh size, then it is obvious that any molecule larger than the size of the mesh will be excluded from the condensate. We would thus expect a hard size cutoff for undesired molecules found in the condensate, as was proposed in [13]. However, biomolecular condensates are typically dynamic network fluids, where local interactions form and dissolve rapidly. It is then unclear how such a liquid condensate could exclude molecules, as it seems large molecules could always sneak into the condensate through large transient holes. In the following, we argue that the conformational entropy of IDRs may be the key factor leading to the exclusion of large particles from condensates. Compared to folded domains, IDRs have much higher conformational entropy, and this implies that removing any of the volume accessible to IDRs has a high free-energy cost. For clients to partition into condensates, they must overcome this cost by binding to scaffolds or other condensate components. Building on this reasoning, we derive a general expression for the resulting entropic exclusion and attractive binding contributions to the partition free energy.

## RESULTS

### Entropy of IDRs excludes particles from a model condensate

To demonstrate the ability of polymer conformational entropy to exclude neutral particles from biomolecular condensates, we performed coarse-grained molecular-dynamics simulations of sticker-spacer polymers [15] (Fig. 2). Briefly, we modeled polymers as linear chains of spherical spacer beads connected by springs. Attached to these spacer beads are sticker beads of two types: A (red) and B (blue). Stickers of different types attract each other via a short-ranged attractive potential, whereas stickers of the same type repel each other to ensure that the heterotypic binding between A and B stickers is one-to-one. The attractive interactions between A and B stickers drive polymer phase separation. Spacer beads (white) and neutral particles (green) interact with each other via short-ranged repulsive interactions; there is no volume exclusion between them and the stickers, i.e., the stickers act as virtual patches to the rest of the system. See Methods for simulation details; all simulations were performed using the LAMMPS Molecular Dynamics Simulator [5, 6].

**FIG. 2.**
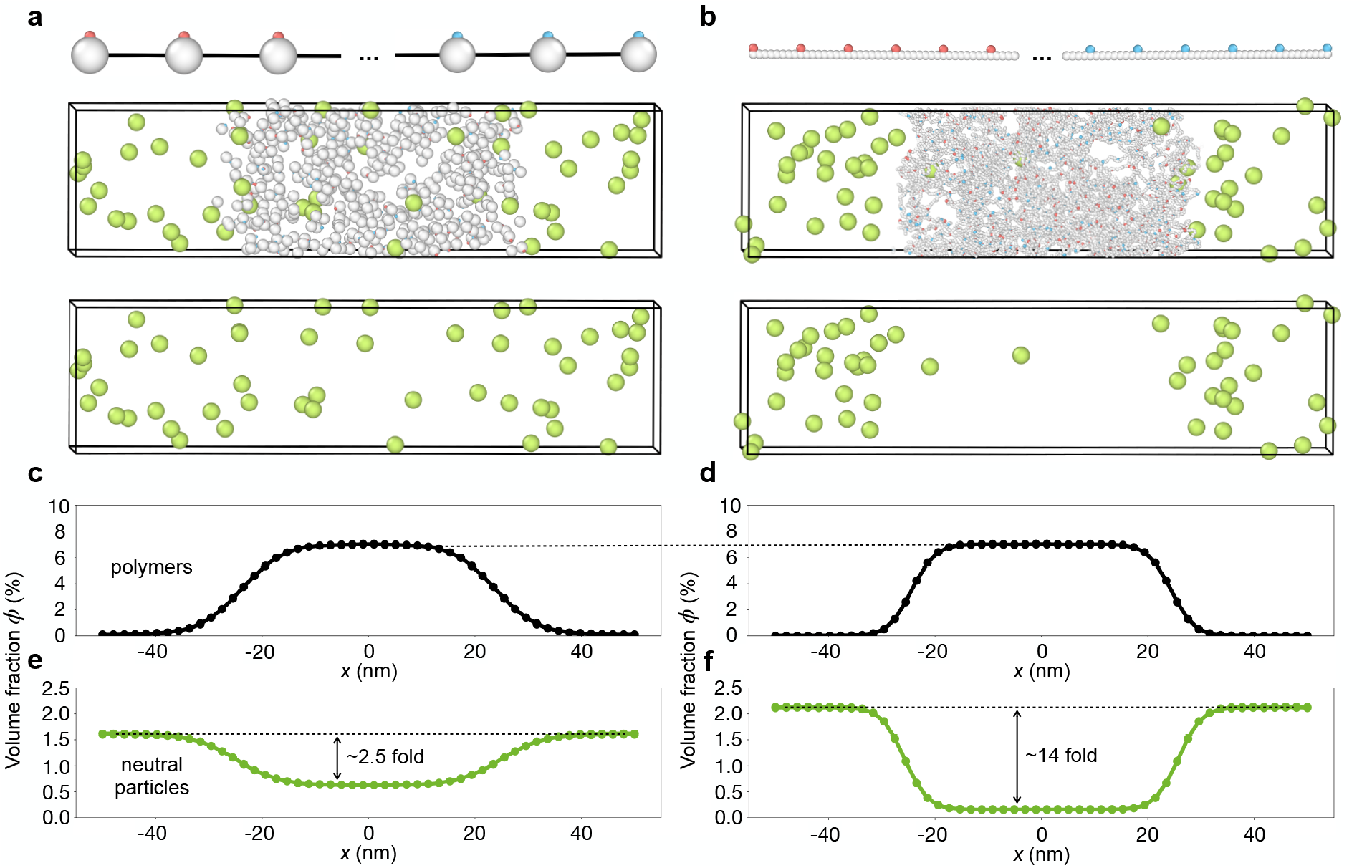
Intrinsically disordered spacer regions strongly exclude neutral particles from a simulated biomolecular condensate. **a**, Snapshot of a coarse-grained molecular-dynamics simulation of 50 sticker-spacer polymers along with 50 neutral particles. Each neutral particle has a radius of 1.5 nm (green). Each polymer is a linear chain of 10 spherical spacers (white) of radius 1 nm (“compact spacers”), connected by stretchable bonds. Each of the 5 spacers at one end of the polymer is attached to a type A sticker (red) and each of the 5 at the other end is attached to a type B sticker (blue). Both types of stickers are of radius 0.3 nm. The system phase separates into a dense phase (middle region) and a dilute phase (two sides) in a 100 nm *×* 25 nm *×* 25 nm box with periodic boundary conditions. **b**, Similar to **a**, but each polymer is a linear chain composed of 450 spherical spacers of radius 0.3 nm (“IDR spacers”), connected by stiff bonds, and with a sticker attached to every 8^th^ spacer along the chain. The total volume of a single polymer equals that of a single polymer in **a. c, d**, Polymer volume-fraction profiles of systems in **a, b** averaged over time and over 60 simulation replicates for **a** and 30 replicates for **b**, with the center of the dense phase aligned at *x* = 0. The volume fraction of the dense phase is 7% in both cases. **e, f**, Neutral particle volume-fraction profiles of systems in **a, b**, respectively. All simulations performed in LAMMPS [5, 6], see Methods for details.

We first simulated a condensate in which the spacers are large spherical beads representing folded modules (“compact spacer”, ∼ 45 amino acids each by volume), with a single small sticker bead attached to each spacer. As shown in Fig. 2a,e, the resulting compact-spacer condensate partially excludes neutral particles, but the concentration of particles is only reduced by ∼ 2.5-fold compared to the surrounding region.

By contrast, as shown in Fig. 2b,f, if we replace each large spacer bead by a string of amino-acid sized beads of the same total volume (“IDR spacer”), with a sticker attached to every 8^th^ bead, the neutral particle concentration in the dense phase is now reduced by ∼ 14-fold. To ensure a fair comparison, we chose parameters such that the polymer volume fraction of the dense phase was the same in both cases, Fig. 2c,d (full phase diagrams are presented in Fig. S1). Thus the only essential difference between the two cases is that the polymers in the IDR-spacer case have more conformational entropy. This simple comparison makes it clear that spacers in the form of IDRs can provide strong exclusion (Fig. 2, right panels), whereas exclusion is nearly absent if the same spacer volume is collapsed into compact domains (Fig. 2, left panels).

### IDRs impose a free-energy barrier to “intruder” particles

The exclusion of neutral particles by explicit IDRs observed in Fig. 2 reflects the free-energy cost of adding a neutral particle to the condensate. In effect, the volume taken up by a neutral particle reduces the possible conformations of the IDRs, and this reduction of IDR entropy implies an increase of the system’s free energy. To quantify this free-energy barrier, we utilized the Widom insertion method [16, 17] to insert a spherical test particle of radius *R* at random locations in the condensates of the types shown in Fig. 2a,b. For each insertion, we calculated the hard-sphere interaction energy *U*(*R*) between the test particle and the local contents of the condensate (see Methods). The average of the resulting Boltzmann factors gives the free-energy cost of insertion:

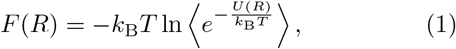

where *k*_B_ is the Boltzmann constant, *T* is the temperature, and the average is over all insertion locations and configurations of the system. As shown in Fig. 3a, over a wide range of test particle radii, the free-energy barrier is substantially higher for the condensate with IDR spacers than for the condensate with compact spacers.

**FIG. 3.**
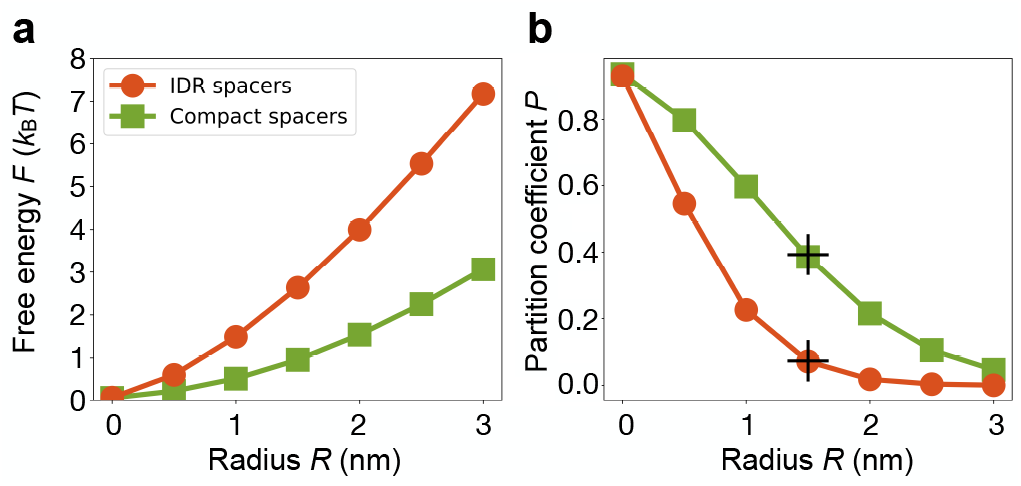
The IDR-spacer condensate excludes neutral particles more strongly than the compact-spacer condensate across a wide range of particle sizes. **a**, Free energy cost *F* of inserting neutral test particles as a function of particle radius *R*, using Eq. (1). Particles are inserted into multiple realizations of the dense phase of the compact-spacer condensate in Fig. 2a (green squares) or the IDR-spacer condensate in Fig. 2b (red circles). **b**, The corresponding partition coefficients *P* of particles predicted by Eq. (2) using Δ*F* = *F*_den_ *− F*_dil_, where *F*_den_ is obtained in **a** and *F*_dil_ = 0 due to the extremely low polymer volume fraction in the dilute phase. The black “+” are data points obtained from the coexistence simulations in Fig. 2e,f, where *P* is the concentration ratio of the 3 nm diameter particles in the dense and dilute phases. See Methods for details.

To understand this difference, consider the probability that a randomly inserted test particle will overlap with a spacer bead in either case. Even though the volume fraction of spacers is the same in both condensates, the probability of an overlap is much higher for the condensate with small spacer beads. This is because the test particle only has to come within the combined radius, *R*_test particle_ + *R*_spacer bead_, to result in a steric overlap. For sufficiently small spacer beads the region of overlap becomes a tube of radius *R*_test particle_ independent of the spacer-bead radius. Thus, in this regime, increasing the number of spacer beads while keeping their total volume fixed increases the length of the tube of “exclusion” without decreasing its radius. This is why the free-energy barrier for test particles in Fig. 3a is always higher for the IDR spacers than for the compact spacers.

The free energy in Eq. (1) yields a prediction for the partition coefficient of particles between dense and dilute phases:

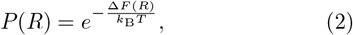

where Δ*F*(*R*) is the free-energy difference between insertion into the dense phase, *F*_den_(*R*), and the dilute phase, *F*_dil_(*R*), both of which can be obtained from Eq. (1). To demonstrate the applicability of this prediction to simulated condensates, we compare the partition coefficient measured directly from coexistence simulations (Fig. 2e,f, shown as the “+” symbols in Fig. 3b) with the values predicted using the Widom insertion method (Fig. 3b). The deviation between measured and predicted partition coefficients is less than 3% for both compact and IDR spacers, verifying Eq. (2).

### Polymer theory for size exclusion

Above, we derived the free-energy barrier in terms of the energetic cost of inserting a test particle, Eq. (1). One can equally derive the free-energy barrier via the reduction of conformational entropy of the IDRs. In a similar vein, it is well-appreciated that maximizing the conformational entropy of polymers is the origin of depletion forces between colloidal particles suspended in a polymer solution [18, 19]. Although such interactions between polymers and particles have been the subject of considerable attention in the polymer physics community [20, 21], there are only a limited number of studies aimed at quantifying the free-energy cost of inserting a particle into a polymer solution [22–25]. These studies suggest that for a semidilute solution of flexible chains in a good solvent, the free-energy cost of inserting a spherical neutral particle obeys the following scaling relationship:

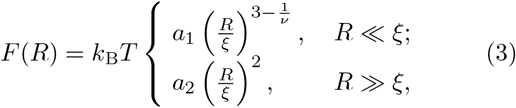

where *R* is the particle radius, *ν* = 0.588 is the Flory exponent for self-avoiding chains, *ξ* is the correlation length of the polymer solution (also known as the “blob size”), and *a*_1_, *a*_2_ are constant prefactors. These expressions describe the free-energy cost for insertion in the two different regimes where the particle radius is either much smaller or much larger than the correlation length. (As discussed below, depending on context the latter regime may also require a term for pressure-volume work.)

To go beyond scaling analysis, we simulated spacers-only polymers held at fixed concentrations as model IDR condensates and quantified the corresponding free-energy cost of neutral particle insertion. Briefly, spacers-only polymers are obtained from IDR-spacer polymers by removing all the stickers. We first simulated these polymers confined in boxes of different sizes to yield varying polymer volume fractions and hence a broad range of correlation lengths *ξ*. The correlation length was calculated using the relationship between the blob size *ξ* and the osmotic pressure Π of the polymer system in the “blob picture”, see Methods:

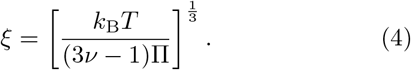

We found that *ξ* ranges from 2.79 nm to 6.10 nm in our systems. We then inserted hard spheres to obtain the free energy cost of insertion (Fig. 4a) using Eq. (1) for each system. Spacers-only polymers require an external pressure Π to maintain a constant volume fraction, which introduces an additional free-energy term, Π(4*πR*^3^/3), for inserted spherical particles of radius *R* due to pressurevolume work. However, for partition into a condensate, this pressure-volume term must be subtracted from the calculated Δ*F*(*R*) which is the free-energy difference for insertion into dense and dilute phases, as the dense and dilute phases are equilibrated at the same pressure. Therefore, in our simulations we evaluated Δ*F*(*R*) by performing Widom insertion into boxes containing self-avoiding polymers under pressure, but then subtracted the product of this pressure and the particle volume.

**FIG. 4.**
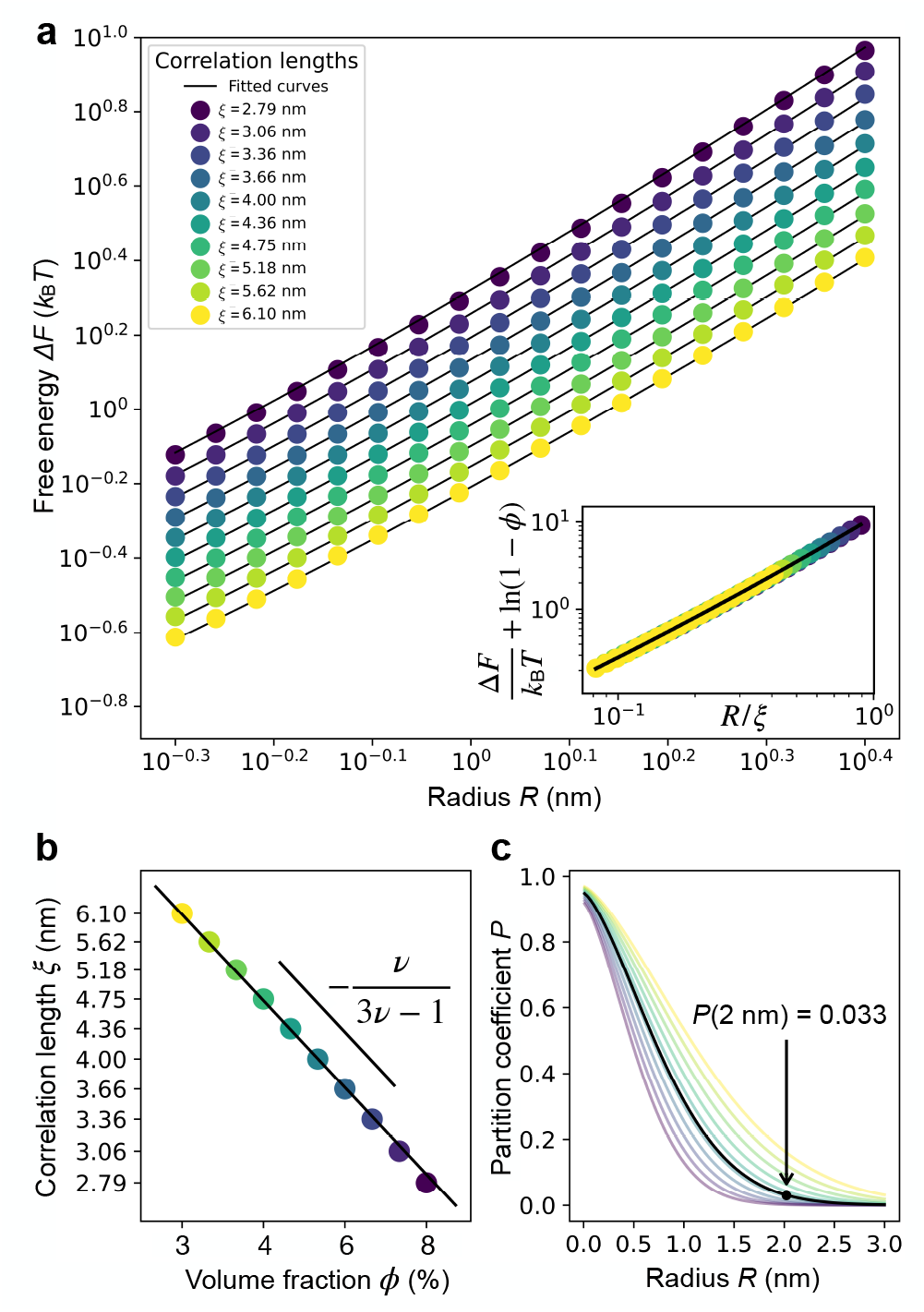
Quantification of the free-energy cost of inserting a hard-sphere particle into homogeneous polymer solutions at various concentrations. **a**, Free-energy cost of insertion as a function of particle radius *R* measured in different simulations using the Widom insertion method, Eq. (1). All simulated polymer systems contain 60 flexible self-avoiding bead-spring polymers each with 450 monomers of radius 0.3 nm (average radius of an amino acid) and equilibrium bond length 0.38 nm (the distance between *C*_*α*_ atoms of consecutive aminoacid residues) in cubic boxes of side length 31.55, 32.72, 33.93, 35.19, 36.49, 37.84, 39.24, 40.69, 42.19 and 43.76 nm, respectively, with periodic boundary conditions. Black curves are from a global fit of the theoretical expression in Eq. (5) to results from all simulations. Inset, data collapse of the inferred polymer-entropy contribution to the adjusted free-energy cost of insertion Δ*F/k*_B_*T* + ln(1 *− ϕ*) as a function of the rescaled particle radius *R/ξ*. **b**, Correlation lengths of the simulated systems generally follow a scaling relationship *ξ∼ ϕ*^*−ν/*(3*ν−*1)^ with *ϕ* the volume fraction of polymers. **c**, Partition coefficient as a function of particle radius for simulated systems. The black curve is for a condensate with a concentration ∼ 250 g/L and a corresponding volume fraction of IDRs ∼ 5% (assume 40% IDR content), predicted by Eqs. (2) and (5) using the correlation length indicated by Eq. (6). Particles as small as 2 nm in radius can be 97% excluded from such a condensate as indicated by the black dot.

To generalize our simulation results, we propose that the free-energy cost of insertion into a condensate can be expressed as

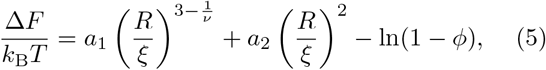

where we have augmented the terms in Eq. (3) with the free-energy cost due purely to volume reduction, *− k*_B_*T* ln(1 *− ϕ*), with *ϕ* being the volume fraction of polymers in the condensate. This term represents the Δ*F*(*R*) for inserting a particle of zero radius, which can be significant for condensates with a high polymer volume fraction. The unified expression, Eq. (5), interpolates between the two limiting scaling behaviors from Eq. (3) while maintaining the correct scaling in each regime. Notably, it implies that if we rescale the radii of inserted particles by the correlation length *ξ* for each polymer system, then Δ*F/k*_B_*T* + ln(1 *− ϕ*) for different systems should collapse onto a universal curve. Indeed, if we rescale the *R* values by the corresponding *ξ* we obtain a collapsed curve (Fig. 4a, inset). Finally, we performed a global fit of all simulation results and obtained the best-fitting parameters *a*_1_ = 4.18 and *a*_2_ = 7.24 (Fig. 4a).

If particles are free to partition from the dilute phase to the polymer-rich region, then according to Eqs. (2) and (5), a particle with a diameter equal to the polymer correlation length *ξ* (i.e., *R* = *ξ/*2) has a free-energy cost of insertion Δ*F* = 3.5 *k*_B_*T* (assuming the free-energy cost of insertion into the dilute phase and the contribution from volume reduction in the dense phase are both negligible). This results in a partition coefficient *P* = 0.03. Thus, the correlation length *ξ* of IDRs in a condensate serves as a strong size cutoff for molecules that lack binding affinity to the condensate.

As a practical matter, the correlation length *ξ* is typically not accessible in experiments. Instead, the dense-phase concentration is the commonly measured quantity from which the volume fraction of IDRs *ϕ* can be estimated. To relate *ξ* and *ϕ*, we note that in polymer physics the two quantities follow a general scaling relationship [26]:

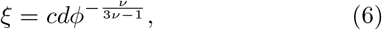

where *d* is the diameter of a monomer and *c* is a constant prefactor. We find that *c* = 0.68 for the fully flexible self-avoiding spacer chains in our simulations (Fig. 4b). Eqs. (2), (5), and (6) can then be utilized to estimate the partition coefficient of a neutral particle of radius *R* in a condensate with a given volume fraction *ϕ*. While in our simulations the volume fractions of polymers in Eq. (5) and of IDRs in Eq. (6) are the same, these quantities will differ for condensates that also contain folded domains. The volume fraction *ϕ* of IDRs in Eq. (6) should then be calculated with respect to the volume available to the IDRs, i.e. not including the volume occupied by folded domains.

Notably, for large intruders, the exclusion free energy in Eq. (5) reduces to

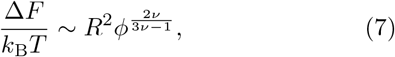

which scales with the surface area of the intruder, rather than its volume. This is because, as remarked above, the condensate and the dilute phase have the same pressure, so there is no pressure-volume work required to move the intruder from one phase to the other. The free energy also increases steeply with the volume fraction of the IDRs, approximately *∼ ϕ*^1.5^.

### Application of the size-exclusion theory to protein condensates

How well does the size-exclusion theory apply to protein condensates? Ref. [8] studied the partitioning of various dextrans into LAF-1 condensates. The measured concentration of LAF-1 protein in the *in vitro* condensates was ∼ 2.6 mM [27]. Each LAF-1 has about 250 amino acids in its disordered regions [28]. Assuming each amino acid is a sphere of radius 0.3 nm, the volume fraction of IDRs is then 3.6%. The IDR correlation length in the LAF-1 condensates according to Eq. (6) (using *c* = 0.68) is then *ξ* = 5.24 nm. The hydrodynamic radii of the 10 kDa, 70 kDa, and 155 kDa dextran probes are 1.9 nm, 6.5 nm, and 8.9 nm, respectively [29]. The free energies of insertion for these dextrans are predicted to be 2.1, 17, and 29 *k*_B_*T* from Eq. (5). Therefore, we expect the 10 kDa dextran to enter the LAF-1 condensate, whereas the 70 kDa and 155 kDa dextrans should be strongly excluded. This is consistent with the observations in [8].

More generally, the scaffold concentrations of biological condensates typically range from 100 to 400 g/L [30]. Consider a typical condensate with a concentration ∼ 250 g/L, if we assume each scaffold molecule has 40% of its amino acids in the IDRs, and take the mass of an amino acid to be 110 g/mole and the radius to be 0.3 nm, then this condensate has a volume fraction of IDRs ∼ 5%, which corresponds to a correlation length of 4.1 nm from Eq. (6) using *c* = 0.68. This suggests that proteins as small as 2 nm in radius (∼ 20 kDa) will already encounter a free-energy cost of 3.4 *k*_B_*T* to enter a typical condensate, yielding a partition coefficient 0.033 (see Fig. 4c). It is therefore likely that exclusion applies universally to the majority of proteins in the proteome, i.e., as long as there is no significant binding affinity between a protein and condensate scaffolds (or clients), the protein is largely excluded. Condensates can then achieve compositional selectivity by recruiting only desired molecules via attractions to condensate components.

### Client recruitment

Biomolecular condensates are selective functional hubs that recruit specific molecules (scaffolds and clients) while excluding others (intruders). The recruitment process is governed by the balance between favorable attraction energies and unfavorable entropic costs of insertion. If one treats these two contributions as independent, the effective free energy for client recruitment can be derived analytically based on the Widom insertion method under reasonable assumptions:

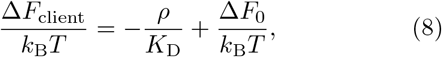

where *ρ* is the concentration of binding sites, each with a dissociation constant *K*_D_ with respect to the clients, and Δ*F*_0_ is the exclusion free energy due to IDRs [Eq. (5)]. In the presence of multiple types of binding sites with different dissociation constants, the first term on the righthand side of the above expression should be summed separately for each type. See Methods for a detailed derivation.

To validate Eq. (8), we introduced attractive interactions between the particles and the stickers in the system shown in Fig. 2b (Fig. 5a). We systematically varied the attraction strength between the particles and the stickers, *ε*, from 1 *k*_B_*T* to 10 *k*_B_*T* in increments of 1 *k*_B_*T*. This range of interaction strengths encompasses the values of *ε* at which the partitioning of particles reverses, transitioning from intruders to clients, Fig. 5b. We next measured the volume fraction of particles from the coexistence simulations and quantified the partition coefficients *P*(*ε*) as the ratio of particles in the dense phase to those in the dilute phase, from which we extracted the recruitment free energy Δ*F*(*ε*)*/k*_B_*T* = *−* ln *P*(*ε*), shown as discrete points in Fig. 5c. To compare the measured free energies with the analytical theory in Eq. (8), we extracted the dissociation constant *K*_D_(*ε*) and the binding site concentration *ρ*(*ε*) from the simulated dense phase for each interaction strength *ε*. The theoretical predictions agree well with the simulation results (Fig. 5c), indicating that Eq. (8) captures the essential physics of client recruitment in our modeled biomolecular condensates. See Methods for details on the theory and simulations.

**FIG. 5.**
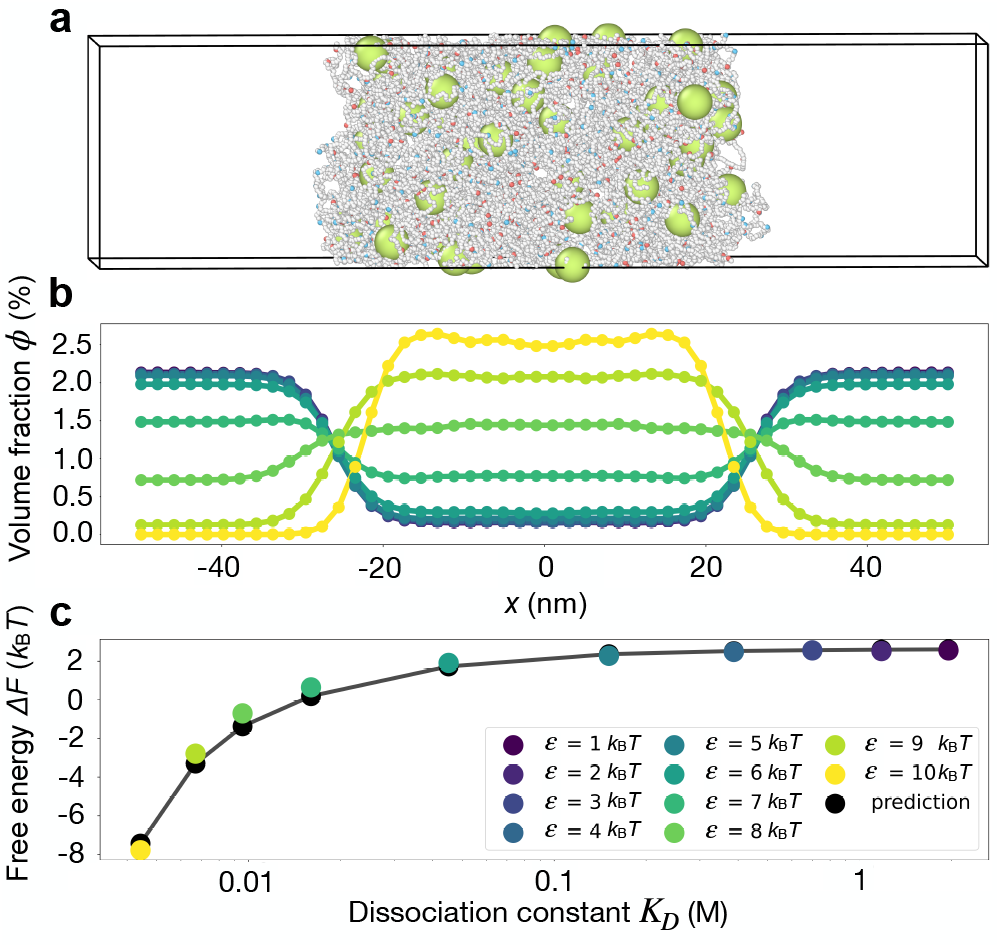
Client recruitment in a modeled biomolecular condensate. **a**, Snapshot of a simulation. The simulation was conducted exactly as in Fig. 2b, except we introduced attractive interactions between the particles (green) and the stickers (red and blue), thereby converting the particles into clients. The attraction strength in this simulation is *ε* = 10 *k*_B_*T*. **b**, Volume fraction profiles of particles in simulated systems with different attractive strengths, *ε* from 1 *k*_B_*T* to 10 *k*_B_*T*. The profiles were obtained by averaging over time and across multiple simulation replicates (10 or 25, depending on *ε*). **c**, Recruitment free energy of the particles inferred from the partition coefficients obtained from **b** (data points), compared to predictions from Eq. (8) (solid curve). See Methods for details.

## DISCUSSION

We presented simulations and theory that support a possible role for protein and RNA IDRs in excluding “unwanted” macromolecules from intracellular condensates.

In a nutshell, IDRs and other flexible polymers enjoy high conformational entropy – thus any restriction of the volume available to these flexible polymers reduces their entropy and so has a free-energy cost. In particular, adding a macromolecule or other particle to a condensate reduces the volume available to the IDRs, which implies the existence of a free-energy barrier to intruders. The magnitude of this barrier depends on the size of the intruding particle as well as on the concentration of the IDRs. Specifically, for large intruders, the barrier scales with the intruder surface area, meaning that larger particles are generally more strongly excluded. That said, inserting a very large neutral particle into a condensate can also incur an energy cost by breaking bonds between stickers [12], making the barrier even higher.

Our proposal that IDRs can exclude unwanted particles from biomolecular condensates is consistent with the many other roles played by IDRs, from holding condensates together, to providing binding sites for clients, to fine-tuning condensate properties. Indeed, one of the proposed roles of IDRs is to reduce condensate density via IDR-IDR self-avoidance [31]. This self-avoidance reflects the same tendency of flexible polymers to maximize their conformational entropy that we explored here as the driving force for exclusion of intruders. Interestingly, the effects of IDR self-avoidance and exclusion could lead to a lower overall volume fraction inside condensates compared to their surroundings in the crowded environment of the cell (see for example Fig. S2).

Going beyond neutral particles, molecules may also be excluded from or enriched in a condensate because of their interactions with the condensate’s chemical environment. It has recently been shown that small molecules, such as metabolites and drugs (*R ∼* 0.5 nm), may preferentially localize inside or outside specific condensates, depending on each molecule’s chemical character [32]. For these small molecules, the free-energy difference in and out of condensates is presumably mostly due to the interactions of the molecules with the chemical environment of the condensate, not to the entropic exclusion from IDRs, as the entropic barrier is negligible for these small molecules (*<* 1 *k*_B_*T*). For larger molecules, such as proteins or RNAs, a combination of entropic exclusion, the chemical environment, and specific interactions with condensate components will contribute to the free-energy of insertion. In the case where the interactions are strong and favorable, the free energy becomes negative, making the molecule a client that preferentially partitions into the condensate.

Looking ahead, we hope that experimental approaches will clarify the extent to which the properties of phase-separating polymers including their IDRs contribute to the exclusion and recruitment of other molecules.

## METHODS

### Simulation details for Figure 2, left panels

We performed coarse-grained molecular-dynamics simulations using LAMMPS [5, 6] to study the exclusion of neutral particles by phase-separated “sticker-spacer” condensates, with compact spacers here and IDR spacers below. Individual polymers in Fig. 2a are modeled as linear chains of 10 spacer beads of type C (white), connected by finitely extensible nonlinear elastic (FENE) bonds

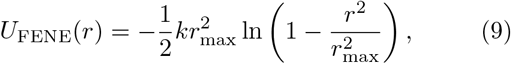

where *k* = 0.129 *k*_B_*T*/nm^2^ is the bond strength constant and *r*_max_ = 14.03 nm is the maximum bond length. These parameters are chosen so that the distance distribution between neighboring spacers is roughly the same as that of the end-to-end distance of 45 smaller spacers in the IDR-spacer case. Spacers have a radius of 1 nm. Each of the first 5 spacers is connected to a sticker bead of type A (red), and each of the following 5 spacers is connected to a sticker bead of type B (blue) via stiff harmonic bonds

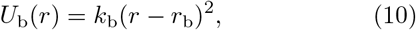

where *k*_b_ = 100 *k*_B_*T*/nm^2^ is the spring constant and *r*_b_ = 1 nm is the equilibrium bond length. Neutral particles of type D (green) are modeled as spherical beads of radius 1.5 nm.

A and B stickers attract each other via a soft potential

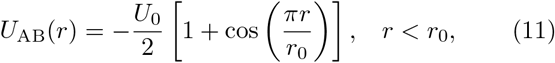

where *U*_0_ = 18.8 *k*_B_*T* is the depth of the potential well and *r*_0_ = 0.3 nm denotes the range of attraction, which equals the radius of a sticker. Various bead types repel each other via a pairwise truncated and shifted LennardJones potential

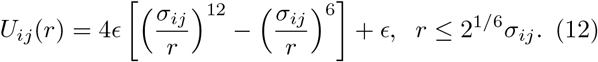

Here, *i* and *j* denote types of particles, where *ij* includes AA, BB, CC, DD, and CD. Note that stickers do not repel from spacers or neutral particles, i.e., they are modeled as virtual patches with respect to the rest of the system; however, they do repel stickers of the same type to avoid many-to-one A:B bindings. *σ* denotes the range of repulsion, *σ*_*ij*_ = *r*_*i*_ + *r*_*j*_, where *r* is the bead radius, *r*_A_ = *r*_B_ = 0.3 nm, *r*_C_ = 1 nm, and *r*_D_ = 1.5 nm. *ϵ* determines the strength of repulsion, which is set to be 1 *k*_B_*T* for all pairs.

We simulated 50 polymers and 50 neutral particles in a 100 nm *×* 25 nm *×* 25 nm box with periodic boundary conditions. We first initialized the simulation by confining polymers in the region 30 nm *< x < −* 30 nm to promote phase separation and ensure that only a single dense condensate is formed. The attractive interaction between A and B stickers [Eq. (11)] was gradually switched on from *U*_0_ = 0 to 18.8 *k*_B_*T* over 4 *×* 10^7^ time steps. This annealing procedure led to the formation of a dense phase close to its equilibrated concentration. The dense condensate was equilibrated at fixed *U*_0_ = 18.8 *k*_B_*T* for another 1 *×* 10^7^ steps and then the confinement walls were removed. The system was equilibrated for 5 *×* 10^7^ more time steps to allow for the formation of a dilute phase and further relaxation of the dense phase. The neutral particles were allowed to equilibrate freely throughout the entire simulation box during these equilibration steps. We finally recorded the positions of all beads every 5 *×* 10^4^ steps for 4000 recordings.

Through the entire simulation, we equilibrated the system using a Langevin thermostat implemented with LAMMPS commands fix nve and fix langevin, i.e., the system evolved according to [33]

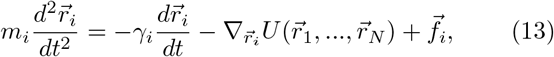

Where 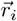 is the coordinate of bead *i, m*_*i*_ is its mass, *γ*_*i*_ is its friction coefficient, 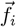 is random thermal noise acting on bead *i*, and the potential energy 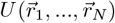 contains interactions between bead *i* and all the rest of beads, including bonds and pairwise interactions [Eqs. (9)–(12)]. We took temperature *T* = 300 K, the damping factor *τ* = *m*_*i*_*/γ*_*i*_ = 1 ns, step size *dt* = 0.01 ns, and mass of particles *m*_A_ = *m*_B_ = 5.65, *m*_C_ = 18.85 ag and *m*_D_ = 28.27 ag. These parameters give each bead the correct diffusion coefficient *D*_*i*_ = *k*_B_*T/*(6*πηr*_*i*_) where *η* is the water viscosity 0.001 kg*/*m*/*s and *r*_*i*_ is the bead radius. 60 simulation replicates were performed with different random seeds to yield robust statistics. Consistency of results was checked across replicates and between the first and second halves of the recorded data.

The volume fraction profiles *ϕ*_C_(*x*) of polymers and *ϕ*_D_(*x*) of neutral particles in Fig. 2c,e were computed along the *x*-axis with a bin size of 1/50 of the box length. For each frame, we identified the center of mass of polymers along *x*-axis and recentered the simulation box to this center of mass. The histograms were averaged over all recordings and simulation replicates, and symmetrized with respect to *x* = 0.

### Simulation details for Figure 2, right panels

The polymers in Fig. 2b represent intrinsically disordered proteins, where each polymer consists of 450 spacers (type C) but with a radius of only 0.3 nm, corresponding to the average radius of an amino acid. The spacers are connected by harmonic bonds with a spring constant *k*_b_ = 692 *k*_B_*T*/nm^2^ and an equilibrium bond length *r*_b_ = 0.38 nm. These parameter choices ensure that each polymer excludes the same volume as the polymer shown in Fig. 2a. Type A stickers are attached to the spacers in the first half of the polymer, while type B stickers are attached in the second half, with one sticker placed on every 8^th^ spacer. These stickers are attached by harmonic bonds with *k*_b_ = 1111 *k*_B_*T*/nm^2^ and *r*_b_ = 0.3 nm. For the interaction between stickers, *U*_0_ was set to 18 *k*_B_*T* in the corresponding soft potential to ensure a similar condensate volume fraction of 7.0% in the two systems shown in Fig. 2a and Fig. 2b.

As in Fig. 2a, we simulated 50 polymers and 50 neutral particles in a 100 nm *×* 25 nm *×* 25 nm box with periodic boundary conditions. Similar to Fig. 2a, the attractive interaction between A and B stickers [Eq. (11)] was gradually switched on from *U*_0_ = 0 to 18 *k*_B_*T* over 4 *×* 10^7^ time steps. The system was equilibrated at fixed *U*_0_ = 18 *k*_B_*T* for another 1 *×* 10^7^ steps, and then the confinement walls were removed. The system was equilibrated for 5 *×* 10^7^ more time steps and then the positions of all beads were recorded every 8 *×* 10^4^ steps for 5000 recordings. We used the Langevin thermostat in Eq. (13) to equilibrate the system. All parameters are the same as in Fig. 2a, except for the mass of particles we took *m*_A_ = *m*_B_ = *m*_C_ = 5.65 ag, and *m*_D_ = 28.27 ag. 30 simulation replicates were performed.

The volume fraction profiles of polymers and neutral particles in Fig. 2d,f were measured following the same procedures as in Fig. 2c,e.

### Derivation of Equation

To derive Eq. (1), we denote by 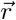 the vector that contains the *x, y, z* coordinates of all polymer beads in the system. The vector 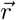 therefore captures the configurations of all polymers. In the canonical ensemble, the probability density *ρ*_0_ for a specific configuration 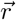 is

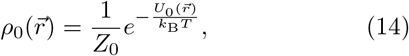

where *Z*_0_ is the partition function of the system

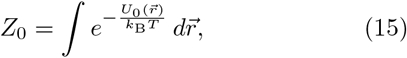

with 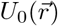 the total interaction energy of a configuration 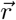. The Helmholtz free energy of the system is then

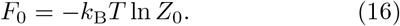

Insertion of a test particle raises the energy of the system. Let 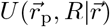 be the potential energy raised by inserting a spherical particle of radius *R* at position 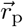 in the polymer configuration 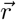. The Helmholtz free energy of the system including the test particle is now

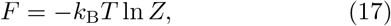

Where

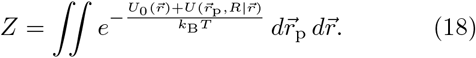

The free energy of insertion (also known as the excess chemical potential) is therefore

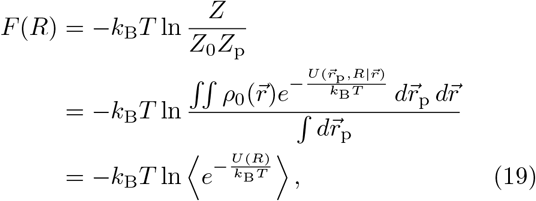

where the average is over all insertion locations and configurations of the system, 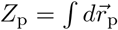 is the reference partition function of the particle for insertion into the same volume but without polymers, and we have abbreviated 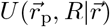 as *U*(*R*).

### Simulation details for Figure 3

We utilized the Widom insertion method, Eq. (1), to quantify the freeenergy cost of inserting a spherical test particle into condensates of the types shown in Fig. 2a,b. Condensate configurations were obtained by recording the positions of all beads every 5 *×* 10^4^ steps for 100 recordings for both types of condensates. We next computed the free-energy costs of inserting spherical particles with radii *R* = 0, 0.5, 1, …, 3 nm into these equilibrated configurations. To ensure that the insertions are within the bulk dense phase, away from the interface, particles were inserted only in the region *−* 10 nm *≤ x ≤* 10 nm, where the polymer volume fraction in the dense phase reaches its plateau value, Fig. 2c,d.

For each configuration, we uniformly sampled 2.744 *×* 10^6^ positions on a 140 *×* 140 *×* 140 grid in the bulk dense phase of the condensate. We note that the test particle only interacts with spacers, as the stickers were modeled as virtual patches. The insertion potential for the hard-sphere particle is therefore:

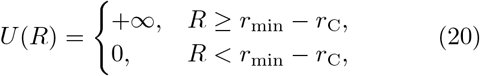

where *r*_min_ is the distance between the particle and the nearest spacer, and *r*_C_ is the spacer radius with *r*_C_ = 1 nm for compact spacers and *r*_C_ = 0.3 nm for IDR spacers, as mentioned earlier. The corresponding Boltzmann factor for each insertion is:

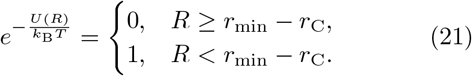

The free-energy cost of insertion and the corresponding partition coefficient can then be calculated according to Eqs. (1) and (2), respectively.

### Derivation of Equation (4)

In the blob picture, polymers in a homogeneous solution spontaneously divide into individual self-avoiding segments or “blobs”. Each blob consists of *g* monomers and behaves as a semi-independent entity within the polymer, i.e., the size of a blob *ξ* scales with *g* as if no neighboring segments/polymers were present

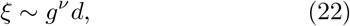

where *ν* is the Flory exponent and *d* is the diameter of a monomer. The number of blobs in a system of total monomers *N* and volume *V* is therefore

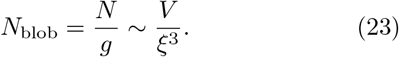

Eqs. (22) and (23) lead to

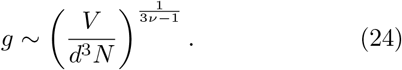

The free energy of the polymer system scales with the number of blobs

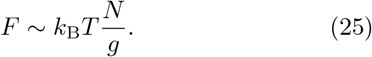

The osmotic pressure is

If we take the prefactor in the last term of Eq. (26) to be 1, we obtain Eq. (4).

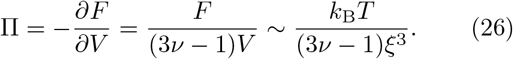

### Simulation details for Figure 4

The free-energy costs of inserting hard-sphere particles into the systems in Fig. 4 were obtained in the following steps. First, we simulated systems with 60 flexible self-avoiding bead-spring polymers each with 450 monomers of radius 0.3 nm (average radius of an amino acid) and equilibrium bond length 0.38 nm (distance between *C*_*α*_ atoms of consecutive amino acids) in, respectively, cubic boxes of side length 31.55, 32.72, 33.93, 35.19, 36.49, 37.84, 39.24, 40.69, 42.19, and 43.76 nm with periodic boundary conditions. Bond potentials follow Eq. (10) with *k*_b_ = 692 *k*_B_*T*/nm^2^ and *r*_b_ = 0.38 nm and pairwise repulsive potentials follow Eq. (12) with ϵ = 1 *k*_B_*T* and *σ* = 0.6 nm. The polymer systems were equilibrated using the Langevin thermostat in Eq. (13) for 10^7^ steps, with temperature *T* = 300 K, damping factor *τ* = 0.01 ns, step size *dt* = 0.0001 ns, and mass *m* = 0.0565 ag. Polymer configurations were recorded every 2 *×* 10^5^ time steps for 2000 recordings. 5 simulation replicates were performed with different random seeds for each system.

Next, the free-energy costs of inserting hard-sphere particles with radii *R* = 0.50, 0.55, 0.60, 0.66, …, and 2.48 nm (18 values linearly spaced on a logarithmic scale) were measured for each system. The insertion procedures are the same as those in Fig. 3, except here for each configuration we uniformly sampled 10^6^ positions on a 100 *×* 100 *×* 100 grid in the boxes. *F*(*R*) of each system was calculated using Eq. (1).

Finally, the osmotic pressures of the polymer systems were measured in LAMMPS using the built-in output variable c thermo press. The pressure was averaged over 4 *×* 10^8^ steps and over all simulation replicates for each system. The measured pressure values are Π = 0.25, 0.19, 0.14, 0.11, 0.085, 0.065, 0.050, 0.040, 0.031, and 0.024 MPa, respectively. We then obtained Δ*F*(*R*) by subtracting the pressure-volume work term Π(4*πR*^3^*/*3) from the above calculated *F*(*R*).

The correlation lengths of the polymer systems in Fig. 4 were found via Eq. (4) using the corresponding osmotic pressures measured above.

### Derivation of Equation (8)

According to the Widom method, the free energy of inserting a test particle with both repulsive and attractive interactions with condensate components can be expressed as:

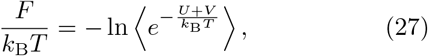

where *U* represents the repulsion from spacers that drives exclusion, while *V* denotes the attraction to binding sites that facilitates recruitment. The average is taken over all insertion locations and configurations of the system. Assuming these contributions are independent, we can decompose the free energy as:

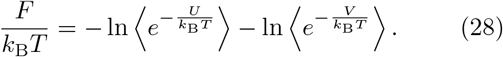

To obtain the attraction free energy, we assume multivalent interactions between the particle and its binding sites, while restricting the maximum number of binding events (valency) to *n*_max_. Thus, if *v* is the interaction potential with a single binding site, we can write:

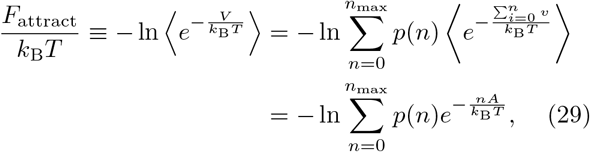

Where

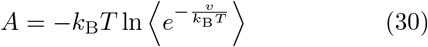

is the effective free energy associated with a single binding site, and *p*(*n*) is the probability that the test particle participates in *n* independent binding interactions, which follows a Poisson distribution:

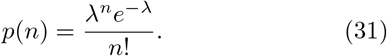

Here, *λ* represents the average number of binding sites with which the test particle interacts in the low binding strength limit. Substituting this expression into the attraction free energy yields:

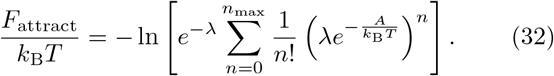

The binding interactions can be characterized by the particle-binding site dissociation constant

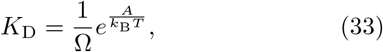

where Ω is the interaction volume between the particle and a single binding site. For a density of binding sites *ρ*, one has *λ* = Ω*ρ*, so that

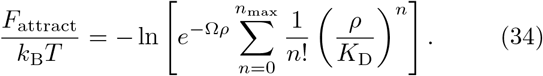

Provided that the typical number of binding interactions is small compared to *n*_max_, the sum becomes an exponential:

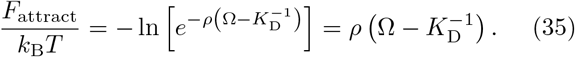

In order for the particle to be recruited, the particle-binding site interaction strength must be much stronger than *k*_B_*T*, therefore *K*_D_ ≪ 1*/*Ω, which simplifies the above expression to:

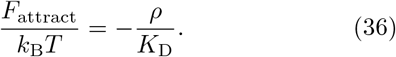

Finally, assuming the free energy of inserting the test particle into the dilute phase is 0, i.e., the dilute phase volume fraction is so low that exclusion is negligible, and there is no strong binding between the test particle and the dilute phase components so that *F*_attract_ there is also negligible, the recruitment free energy, Δ*F* = *F*_den_ *− F*_dil_, then takes a general form:

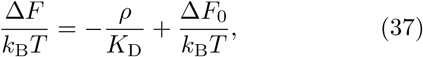

which is Eq. (8), where Δ*F*_0_ is the exclusion free energy in Eq. (5). The above result naturally generalizes to systems with multiple types of independent binding sites:

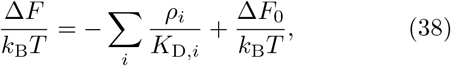

where *ρ*_*i*_ represents the concentration of binding site type *i*, and *K*_D,*i*_ is the corresponding dissociation constant.

### Simulation details for Figure 5

The condensate characteristics in Fig. 5 match those depicted in Fig. 2b. To model client recruitment, we introduced an attractive potential between type D particles and both sticker types (A and B), defined as:

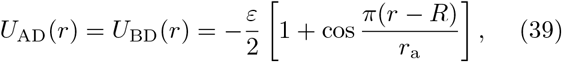

for *R − r*_a_ *≤ r ≤ R* + *r*_a_; *U*_AD_(*r*) = *U*_BD_(*r*) = 0, otherwise. This potential establishes a smooth, short-range attraction between a particle and a sticker. We employed 10 values for the depth of the potential well, *ε*, ranging from 1 *k*_B_*T* to 10 *k*_B_*T*. The particle radius was set to be *R* = 1.5 nm and the attraction range *r*_a_ = 0.3 nm.

Simulation parameters and procedures were adopted from those used in Fig. 2b. A total of 2.5 *×* 10^8^ steps were performed after equilibration for all *ε* values. Volume fraction profiles were obtained in the same way as in Fig. 2e,f. For *ε* values in the range from 1 *k*_B_*T* to 5 *k*_B_*T*, 10 replicas sufficed for a smooth profile. Simulations with *ε* in the range from 6 *k*_B_*T* to 10 *k*_B_*T* required 25 replicas due to the reduced sampling efficiency. Partition coefficients, *P*(*R, ε*), were calculated as

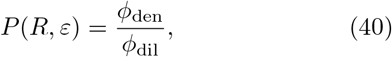

where the dense-phase volume fraction was averaged over the range *−* 10 nm *< x <* 10 nm, and the dilute-phase volume fraction was averaged over *x <* − 40 nm and *x > −* 40 nm. The recruitment free energy was then determined for each value of *ε* as:

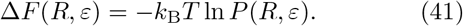

To test the validity of Eq. (8), we obtained the value of *K*_D_ directly from simulations for each choice of *ε*. By definition

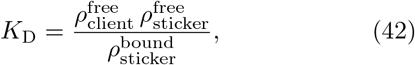

where the *ρ*’s indicate the relevant densities in the dense phase after equilibrium. Note that we used *ρ*^free^ *ρ*_client_, which is the total client density, both free and bound. This is because in our simulations a client can interact with multiple stickers so all clients are effectively “free”. We obtained *K*_D_ directly from simulations using Eq. (42).

## ACKNOWLEDGMENTS

VG and YZ were supported by a startup fund at Johns Hopkins University. NSW was supported by the National Science Foundation, through the Center for the Physics of Biological Function (PHY-1734030), NIH Grant R01 GM140032, and the Princeton Biomolecular Condensate Program. This work was carried out at the Advanced Research Computing at Hopkins (ARCH) core facility (rockfish.jhu.edu), which is supported by the National Science Foundation (NSF) grant number OAC1920103.

