## Supplemental Material for "Conformational entropy of intrinsically disordered proteins bars intruders from biomolecular condensates"

### SUPPLEMENTAL FIGURES

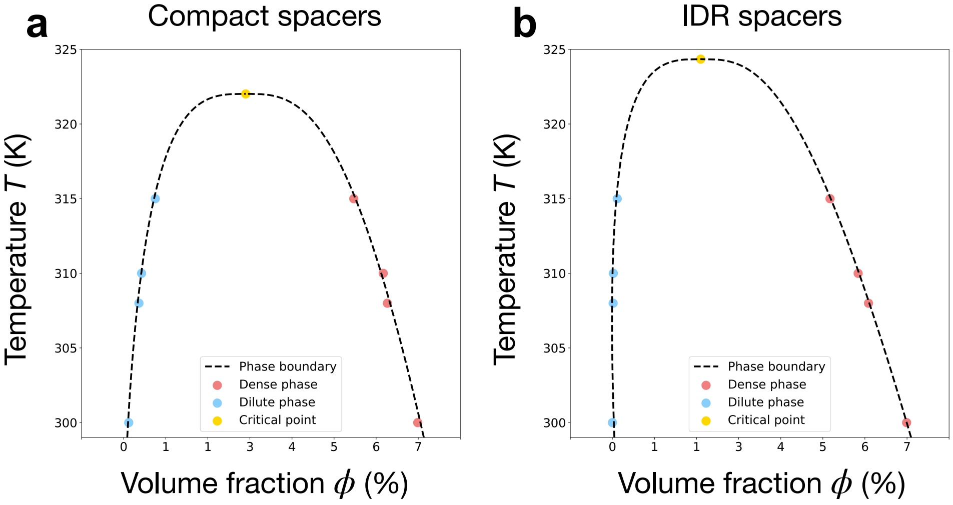


FIG. S1. Phase diagrams for the sticker-spacer polymer systems, **a** for compact spacers shown in Fig. 2a and **b** for IDR spacers in Fig. 2b. 10 replicas were simulated at each choice of temperature for both systems. The volume fractions of polymers were averaged over $-10\,\text{nm} < x < 10\,\text{nm}$ to yield $\phi_{\text{den}}$ and over $x < -40\,\text{nm}$ and $x > 40\,\text{nm}$ to yield $\phi_{\text{dil}}$. The critical points and phase boundaries were determined using the law of coexistence densities for liquid-vapor coexistence $[\phi_{\text{den}}(T) - \phi_{\text{dil}}(T)]^{3.06} = d(1 - T/T_{\text{c}})$ and the law of rectilinear diameters $[\phi_{\text{den}}(T) + \phi_{\text{dil}}(T)]/2 = \phi_{\text{c}} + A(T - T_{\text{c}})$ by fitting $d$ and $A$.

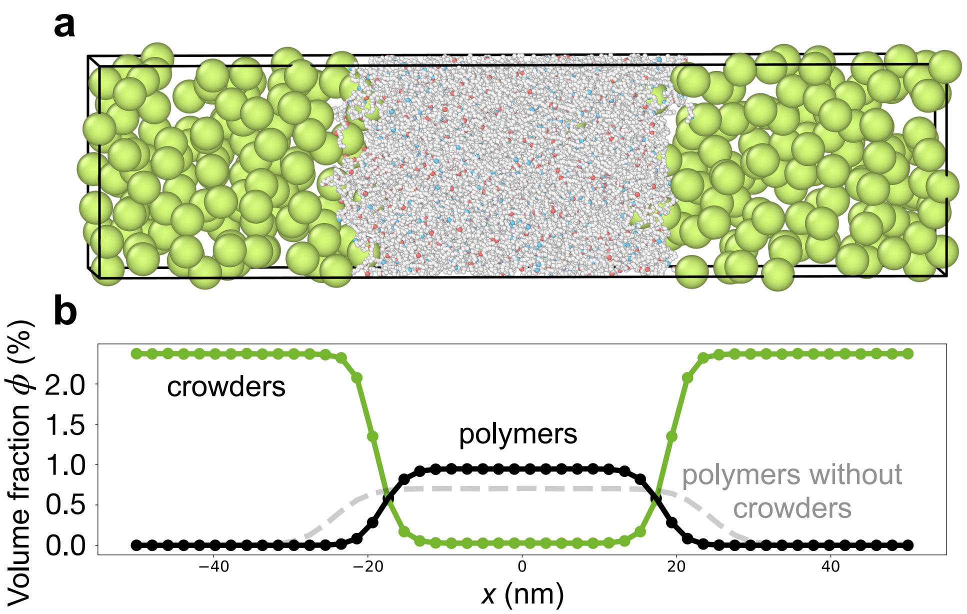


FIG. S2. Demonstration of the crowding effect on a simulated biomolecular condensate. **a**, Snapshot of the simulation. **b**, Volume fraction profiles of polymers (black) and particles (green) in the simulated system. The simulation was performed exactly as in Fig. 2b but with a radius of 2.0 nm for the green particles (the average radius of a globular protein, as reported in [1]) and an increased number of green particles (280) to match the physiological crowder volume fraction in the cytoplasm, which is 20–30% [2]. The gray curve is the volume fraction profile of polymers after removing the particles from the system. While the presence of crowders increases the overall occupied volume fraction in the dense phase, the volume fraction in the surrounding dilute phase is higher still. This follows from the pressure balance between the two phases: the high conformational entropy and self-avoidance of the IDRs in the condensate create a high pressure, which has to be matched by an even higher volume fraction of crowders in the dilute phase.

[1] V. P. Zhdanov, Conditions of appreciable influence of microrna on a large number of target mrnas, Molecular bioSystems **5**, 638 (2009).

[2] J. S. Kim and A. Yethiraj, Effect of macromolecular crowding on reaction rates: a computational and theoretical study, Biophysical journal **96**, 1333 (2009).
